# Oestrogen influences B cell class-switching in individuals with an XX sex chromosome complement

**DOI:** 10.1101/2024.07.18.604160

**Authors:** Hannah Peckham, Anna Radziszewska, Nina M de Gruijter, Restuadi Restuadi, Melissa Kartawinata, George A Robinson, Lucia Martin-Gutierrez, Claire T Deakin, Lucy R Wedderburn, Elizabeth C Jury, Gary Butler, Elizabeth C. Rosser, Coziana Ciurtin

**Affiliations:** Centre for Adolescent Rheumatology Versus Arthritis at UCL, UCLH and GOSH, London, UK; Centre for Rheumatology, University College London, London, UK; Infection, Immunity and Inflammation Research & Teaching Department – UCL Great Ormond Street Institute of Child Health, London, UK; University College London Hospital, London, UK; NIHR Biomedical Research Centre at Great Ormond Street Hospital, London, UK; School of Population Health, Faculty of Medicine and Health, University of New South Wales, Australia

## Abstract

Sex differences in humoral immunity are well-documented, though the mechanisms underpinning these differences remain ill-defined. Here, we demonstrate that post-pubertal cisgender females have higher levels of class-switched B cells compared to age-matched cisgender males. However, whilst sex chromosome-encoded genes characterise most of the differences in total B cell transcriptomes between cisgender-females and -males, sex differences in class-switched B cells are only observed post-pubertally. Accordingly, B cells express high levels of oestrogen receptor 2 *(ESR2)* and genes known to regulate B cell class-switching are enriched for *ESR2-*binding sites. Using a gender-diverse cohort of young people, we show that in transgender males (XX chromosomal background), blockade of natal oestrogen reduced the frequency of class-switched B cells, whilst gender-affirming oestradiol treatment in transgender females (XY chromosomal background), did not increase the frequency of class-switched B cells. These data demonstrate that sex hormones and chromosomes work in tandem to impact immune responses, with oestrogen only supporting B cell class-switching on an XX chromosomal background.

**eTOC summary:** Sex hormones and chromosomes work in tandem to impact immune responses, with oestrogen influencing B cell class-switching exclusively on an XX chromosomal background.

## INTRODUCTION

The presence of a sex bias in the human immune system has been widely reported in recent decades. Studies from across the field of infection and immunity highlight the need to disaggregate research findings by sex and gender. Typically, cisgender males (cis-males) appear more at risk from many infectious diseases, as evidenced by the recent Coronavirus disease 2019 (COVID-19) pandemic (Klein *et al*., 2020; Peckham *et al*., 2020; Scully *et al*., 2020; Biswas *et al*., 2021; Takahashi and Iwasaki, 2021), while cisgender females (cis-females) are more likely to develop humoral autoimmune diseases such as systemic lupus erythematous (SLE) (Lim *et al*., 2014; Brinks *et al*., 2016; Izmirly *et al*., 2017; Conrad *et al*., 2023) Sjögren’s disease (Beydon *et al*., 2024), dermatomyositis (Phillippi *et al*., 2017; Kronzer *et al*., 2023) and systemic sclerosis (Ciaffi et al., 2021; Hughes et al., 2020; Zhong et al., 2019). Very little is known regarding immune-mediated disease outcomes in transgender females and males (trans-females and -males) or non-binary people (those identifying neither as exclusively male nor exclusively female), in the context of gender-affirming hormone therapy (GAHT) treatment (Peckham *et al*., 2022). Studies that aim to investigate the immunological underpinnings of sex and gender biases in inflammatory disease remain of critical importance.

Socioeconomic determinants relating to differing gender roles are known to play a part in disease risk and outcomes (Morgan and Klein, 2019; Bischof *et al*., 2020). However, there is strong evidence that sex chromosomes (X and Y) and sex hormones are the driving influences behind sex biases in conditions affecting the immune system. The X chromosome, of which cis-females and trans-males have two copies, encodes the most immune-relevant genes from the human genome (Bianchi *et al*., 2012). X chromosome inactivation (XCI) silences the expression of one copy of each X gene (in people with two X chromosomes), but a subset of genes may exhibit variable tissue-specific differences in XCI. Key examples include immune-related genes such as toll-like receptor-7 (*TLR7*), which is well-documented in its relevance for immune signalling and implications for autoimmunity (Qu *et al*., 2015; Wang *et al*., 2016; Tukiainen *et al*., 2017; Souyris *et al*., 2018; Oghumu *et al*., 2019); CD40 ligand (*CD40-L)*, vital to T and B cell activation signalling that maintains immune tolerance (Pucino, Gardner and Fisher, 2020); Nuclear factor kappa-light-chain-enhancer of activated B cells (*NF-κB),* which is involved in B cell fate decisions throughout the immune response (Guldenpfennig, Teixeiro and Daniels, 2023); the interleukin-2 (IL-2) receptor gamma chain (*IL-2RG*), mutations of which lead to X-linked severe combined immunodeficiency (SCID) (Tuovinen *et al*., 2020); and Bruton’s tyrosine kinase (*BTK*) (Hagen *et al*., 2020), which regulates B cell activation and differentiation. XCI is triggered on the future inactive X (Xi) by the build-up of *XIST* long coding RNA (Loda, Collombet and Heard, 2022), which recruits a program of gene-silencing factors. By perturbing the expression of *Xist* in a mouse model, a recent study proposed a direct link between dysregulated XCI and the development of autoimmunity (Huret *et al*., 2023). Furthermore, males with Klinefelter syndrome, who have one or more additional X chromosomes, have an increased risk of developing humoral autoimmune disorders such as SLE (Scofield *et al*., 2008; Dillon *et al*., 2011), systemic sclerosis (Rovenský *et al*., 2014) and Sjögren’s Disease (V. M. Harris et al., 2016), despite the presence of male sex hormones.

However, sex biases in immune-mediated diseases are not consistent throughout age. Disorders such as asthma and atopy, characterised by male preponderance pre-puberty, have increased female prevalence during reproductive years (Osman, 2003). In addition, the onset of various adolescent autoimmune conditions at the time of puberty (Whincup *et al*., 2001; Cattalini *et al*., 2019; Massias *et al*., 2020) suggests that increased endogenous exposure to sex hormones, as well as age, have a direct impact on immune system function (Nikolich-Žugich, 2017; Sadighi Akha, 2018). There is a growing body of evidence to suggest that sex hormones influence the development of autoimmunity (McMurray and May, 2003; Petri, 2008; Kim *et al*., 2022), allergy and asthma (McCleary *et al*., 2018; Han, Forno and Celedón, 2020) and the effectiveness of pathogen clearance (Gupta et al., 2022; Ruggieri et al., 2018. The principal sex hormones in humans are testosterone (an androgen), progesterone (“P4”, a progestin), and the four major forms of oestrogen (E1/2/3/4). Intracellular receptors for these hormones are found on numerous cell types, including many immune cells (*The human protein atlas*, 2000). For oestrogen, the main receptors in humans are Oestrogen Receptor alpha (ERα/ESR1) and Oestrogen Receptor beta (ERβ/ESR2), while the Androgen Receptor (AR) is the main receptor for testosterone. These receptors act as key transcriptional regulators throughout the body, with ligation by their cognate hormone impacting both physiological, metabolic, and immune system function (Gegenhuber *et al*., 2022; Hayakawa *et al*., 2022; Lucas-Herald and Touyz, 2022; Hoffmann *et al*., 2023).

Studies comparing the immune system between females and males have long established that there are sex-based differences in B cell responses, with females displaying higher basal and post-vaccination antibody titres than males (Flanagan *et al*., 2017). Studies in humans have suggested that this is due to the suppressive effect of testosterone (Furman *et al*., 2014). However, murine literature suggests that oestrogen pays a dominant role in regulating class-switch recombination (CSR), the process by which B cell receptors switch the constant region of their immunoglobulin from initial IgM and IgD isotypes to IgG, IgA or IgE. IgG antibodies can pass from the blood into tissue sites, and thus have particular importance in viral clearance and immunisation response. In the context of autoimmunity, IgG antibodies also make up the bulk of the pathogenic *auto*antibodies (Lin and Li, 2009; Villalta *et al*., 2013; Olsen and Karp, 2014; Jayakanthan *et al*., 2016), being found in the kidney biopsies of patients with SLE nephritis (Bijl, 2002; Kenderov *et al*., 2002) and demonstrated as damage-causing in mouse models (Madaio *et al*., 1987; Pankewycz, Migliorini and Madaio, 1987). Thus, heightened IgG production may also contribute to the increased risk of autoimmune development in females. Conversely, autoantibodies of the IgG subclass IgG4 have been shown to have protective or anti-inflammatory effects (Pan *et al*., 2020; Jiang *et al*., 2022). The molecular mechanism, as well as the relative contribution of sex hormones and sex chromosomes to B cell class-switching in humans remains ill-defined.

Here, we have utilised a unique gender-diverse cohort of pre-pubertal and post-pubertal cisgender and post-pubertal transgender healthy young people to investigate the relative contribution of sex hormones and sex chromosomes in driving differences within the immune, and more specifically humoral immune, system. Collectively, our data demonstrate that female sex chromosomes and sex hormones work in tandem to influence B cell class-switching in humans. Due to the important role that B cells have in controlling multiple immune-mediated disease outcomes, these data provide significant insights into the clinical sex biases observed in these scenarios.

## RESULTS & DISCUSSION

### Post-pubertal cisgender females have an increase in class-switched memory B cells within peripheral blood when compared to post-pubertal cisgender males

To interrogate which cell population/s may be behind the observed sex differences in immunity, the proportion of 31 innate and adaptive immune cell types within peripheral blood samples were compared between 103 post-pubertal cisgender healthy females (cis-females) and males (cis-males) using flow cytometry (**Figure 1A and Table 1 for demographic information)**. Multiple immune cell frequencies were differentially expressed based on sex, including natural killer (NK) cells, CD4^+^ T regulatory cells (Tregs), total CD8+ T cells, naïve CD8^+^ T cells, naïve (CD27^-^IgD^+^) B cells and ‘double negative’ (CD27^-^IgD^-^) B cells; however, CD19^+^CD27^+^IgD^-^ class-switched memory B cells displayed the most striking sex differences (**Figure 1A-F**). More specifically, cis-females had significantly elevated relative proportions of CD19^+^CD27^+^IgD^-^ class-switched memory B cells compared to cis-males (**Figure 1C**) which were associated with an increase in IgG^+^ B cells in cis-females compared to cis-males with no significant differences in the levels of IgA^+^ or IgE^+^ B cells between the sexes (**Figure 1G-H** and **Supplementary Figure 1A-D**). This corroborated previous studies which have shown increased IgG titres in females following vaccination (Fink et al., 2018).

**Figure 1:**
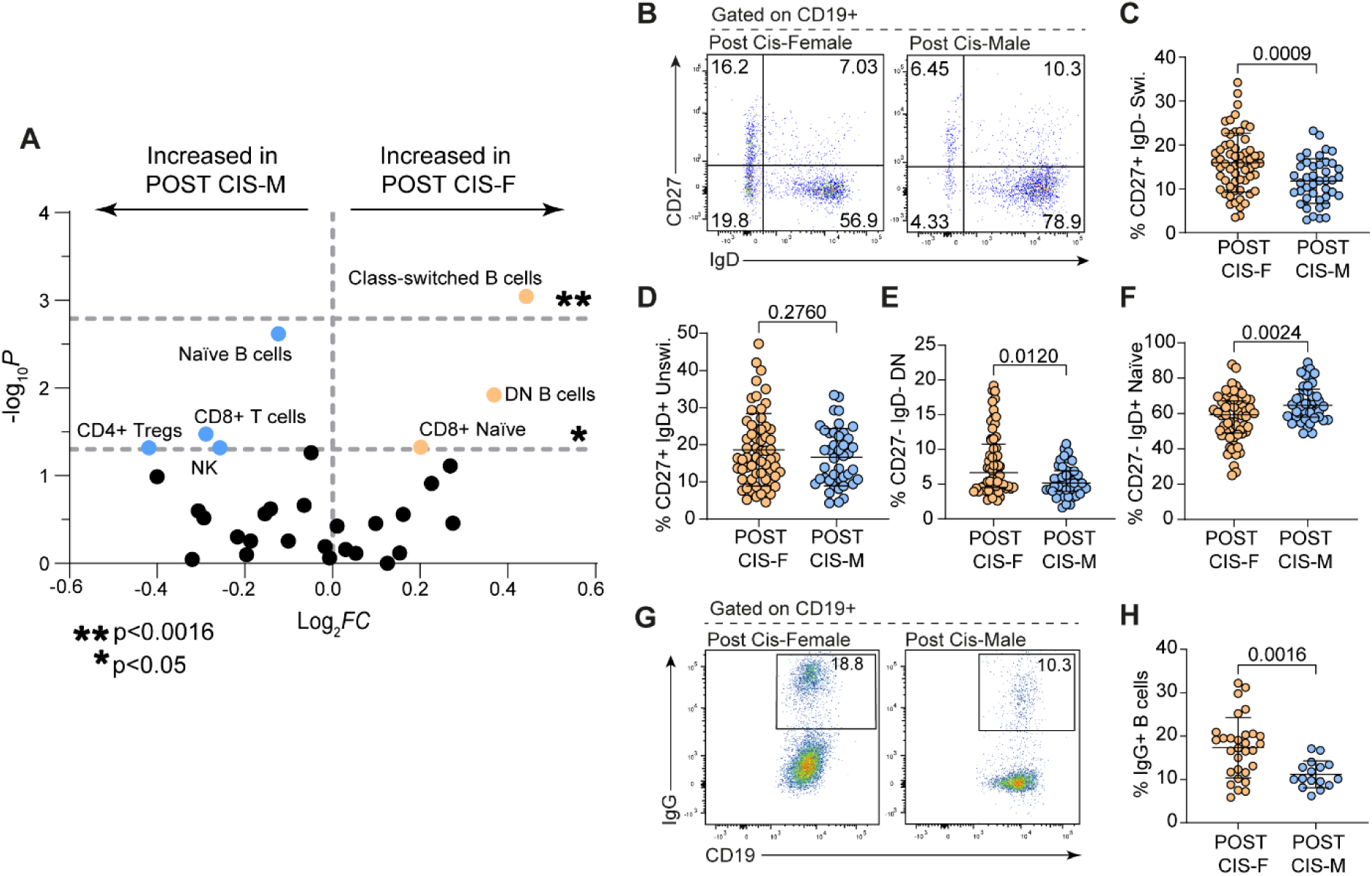
Healthy post-pubertal cisgender females have a higher percentage of class-switched B cells than age-matched healthy cisgender males. (**A**) Volcano plot showing - log_10_ p-values and log_2_ fold changes of flow cytometric relative percentages of peripheral blood immune cell subtypes between post-pubertal cisgender female (POST CIS-F) and male (POST CIS-M) healthy donors (**horizontal dashed lines show p<0.05 and multiple testing adjusted significance level of p<0.0016)**. (**B)** Representative flow cytometry plots and summary dot plots comparing POST CIS-F and POST CIS-M percentages of **(C)** Class-switched memory (CD27+ IgD-), **(D)** Unswitched (CD27+ IgD+), **(E)** Double Negative (CD27-IgD-) and **(F)** Naïve (CD27-IgD+) B cells (POST CIS-F n=61; POST CIS-M n=42). **(G-H)** Representative flow cytometry plots and summary dot plots comparing POST CIS-F and POST CIS-M percentages of IgG+ B cells (POST CIS-F n=30; POST CIS-M n=17). Unpaired t-test with Mean +SD (C, D, H) or Mann-Whitney U-test with Median +IQR (E-F) as appropriate to distribution of data.

**Table 1:** Demographic information for cisgender (pre-pubertal and post-pubertal) and transgender healthy control sample donors used for the immunophenotyping cohort. HC-healthy control; GnRHa-Gonadotrophin releasing hormone agonists; IM-Intramuscular. Fisher’s exact test used to calculate differences between groups.

| **Immunophenotyping HC Cohort** | **Pre-Pubertal Cis HC** | **Post-Pubertal Cis HC** | **Trans HC** | **P-value**<br>**a) Pre vs. Post**<br>**b) Post vs. Trans** |
| --- | --- | --- | --- | --- |
| **Total n** | 26 | 103 | 80 | - |
| **Gender Identity F:M** | 15:11 | 61:42 | 33:47 | - |
| **Mean Age- years (range)** | 8.5<br>(6-11) | 19.9<br>(14-32) | 18.0<br>(15-19) | a) N/A<br>b) 0.0587 |
| **Tanner Stage 1 (Pre-Puberty) (%)** | 26 (100) | 0 (0) | 0 (0) | - |
| **Tanner Stage 2-3 (In Puberty) (%)** | 0 (0) | 0 (0) | 2 (2.5) | - |
| **Tanner Stage 4-5 (Completing/Completed Puberty) (%)** | 0 (0) | 103 (100) | 78 (97.5) | - |
| **Mean Age of Menarche- (years) in XX individuals (range)** | N/A | 12.5 (9-17) NB data on 58/61 donors | 12.1 (9-14) | a) N/A<br>b) 0.2275 |
| **Ethnicity (%)** |  |  |  |  |
| **Asian (any)** | 2 (7.7) | 27 (26.2) | 1 (1.3) | a) 0.0632<br>b) <0.0001 |
| **Black (any)** | 0 (0) | 9 (8.7) | 1 (1.3) | a) 0.2027<br>b) 0.0445 |
| **Other/Mixed** | 2 (7.7) | 8 (7.8) | 6 (7.5) | a) >0.9999<br>b) >0.9999 |
| **White (any)** | 22 (84.6) | 59 (57.3) | 72 (90) | a) 0.0119<br>b) <0.0001 |
| **Medications** |  |  |  |  |
| **Oral Contraceptive (%)** | 0 (0) | 10 (9.7) | 0 (0) | - |
| **GnRHa “Blockers” F:M (%)** | N/A | N/A | 33:40<br>(91.3) | - |
| **Oestradiol (trans-f; oral/patch) (%)** | N/A | N/A | 20 (25) | - |
| **Testosterone (trans-m; IM) (%)** | N/A | N/A | 25 (31.3) | - |
| **Mean months on oestradiol (trans-f) (range)** | N/A | N/A | 13.3 (4.5-23) | - |
| **Mean months on testosterone (trans-m) (range)** | N/A | N/A | 14.6 (4-43) |  |

### Sex differences in the B cell transcriptome between healthy, post-pubertal cisgender females and males are driven by sex chromosome-encoded genes

To understand the potential mechanistic underpinnings of the expansion of CD19^+^CD27^+^IgD^-^ class-switched memory B cells in cis-females compared to cis-males, we next analysed the transcriptome of total CD19^+^ B cells from post-pubertal cis-females and post-pubertal cis-males via RNA sequencing (RNAseq). The transcriptome of B cells was clearly segregated by sex (**Figure 2A**) with 64 differentially expressed genes (DEGs; full gene list with chromosome number/letter shown in **Supplementary Table 1**) between these two groups, of which 28 were downregulated in cis-females compared to cis-males, and 36 were upregulated. 40 (62.5%) of the identified DEGs were found to be encoded on either the X (22/40) or Y (18/40) chromosome (**Figure 2B-C** and **Supplementary Table 1**). As expected, this included genes such as *XIST* and *TSIX*; long non-coding RNA that control silencing of the X chromosomes. The level of expression of *XIST* and *TSIX* varied amongst cisgender females, which could have implications regarding differences in the extent of XCI amongst individuals. In terms of B cell biology, dysregulation in *XIST* has been recently shown to lead to the expansion in the CD11c+ atypical B cells in female patients with SLE or COVID-19 (Yu *et al*., 2021). However, we did not find any correlation between the levels of class-switched memory B cells and *XIST* transcript levels in our cohort (data not shown). Of the DEGs that were not X or Y genes, few had clear known immune functions in B cells with the exception of *HRK* and *CCL5* which were downregulated in cis-females and *IRAK3* which was upregulated in cis-females. Importantly, Harakiri (*HRK)* is in the BCL2 family of proteins which regulates cell death (Inohara *et al*., 1997; Kaya-Aksoy *et al*., 2019; Spetz, Presser and Sarosiek, 2019), CCL5 is a known chemoattract (Aldinucci, Borghese and Casagrande, 2020; Gauthier *et al*., 2023) and IRAK3 is a negative regulator of TLR signalling (Pereira and Gazzinelli, 2023). Further studies are needed to interrogate the function of these genes in B cell function to understand their impact on any of our observed sex differences in the B cell compartment. Pathway analysis of DEGs upregulated in cis-females compared to cis-males revealed significant enrichment of gene ontology biological pathways (GO BP) entitled *translational initiation, negative regulation of protein catabolic process, ribosome biogenesis* and *establishment of protein localisation to organelle*, whilst pathway analysis of DEGs downregulated in cis-females compared to cis-males revealed significant enrichment of GO BP pathways entitled *translation, secretion by cell and cellular response to growth factor stimulation* (**Figure 2D and E**). Changes in these pathways further support a role for sexually divergent B cell activation, differentiation and potentially class-switching profiles, as B cell maturation is well-known to be associated with drastic alterations in cellular structure and translational machinery needed to produce vast amounts of protein in the form of immunoglobulin (Nutt *et al*., 2015).

**Figure 2:**
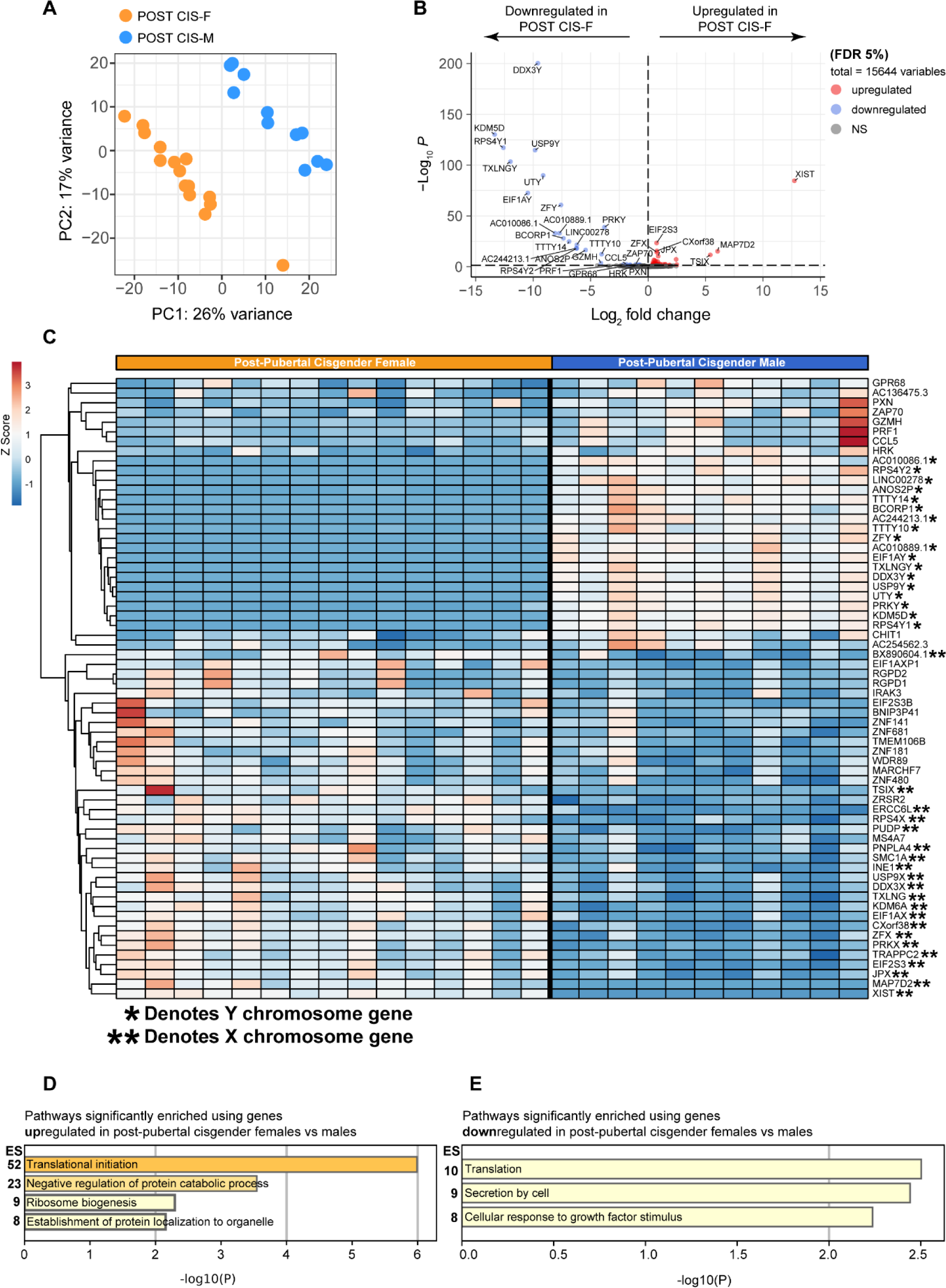
Sex chromosome genes dominate the differential gene expression between total CD19+ B cell RNA from healthy, young cisgender females and males. (**A**) PCA plot showing clustering of post-pubertal cisgender female (POST CIS-F) vs. post-pubertal cisgender male (POST CIS-M) CD19+ B cell gene expression. **(B)** Volcano plot shows differentially expressed genes that are up-(red) or down-(blue) regulated in POST CIS-F compared to POST CIS-M. Only those with a padj<0.05 are labelled as significant. **(C)** Heatmap demonstrating clustering by DEG profiles of POST CIS-F vs. POST CIS-M. **(D-E)** Metascape pathway enrichment analysis (PEA) shows pathways significantly enriched in analysis of **(D)** upregulated DEGs and **(E)** downregulated DEGs in POST CIS-F compared to POST CIS-M (n=15 POST CIS-F, 11 POST CIS-M). ES-‘Enrichment Score’ (the ratio of the proportion of genes in the list that were associated with the enriched term to the proportion of genes in the genome that are associated with the enriched term)

### Sex differences in B cell class-switching are not observed between pre-pubertal cisgender females and males

If the sex chromosomes are, indeed, solely responsible for the observed sex differences in B cell class-switching, it may be expected that these differences would be observed across all age brackets despite changing levels of sex hormones. However, when comparing the frequency of class-switched B cells in healthy, pre-pubertal cisgender females and males (where levels of endogenous sex hormones are very low), the proportions of CD19^+^CD27^+^IgD^-^ memory B cells and of IgG^+^ class-switched B cells were not significantly different between pre-pubertal cis-females (referred to as PRE CIS-F in figures) and cis-males (referred to as PRE CIS-M in figures) (**Figure 3A-E and Supplementary Figure 1A-D**). Furthermore, when each pre-pubertal sex was compared to their post-pubertal sex-matched counterparts, significantly increased proportions of total class-switched B cells were observed in post-pubertal versus pre-pubertal cis-females (**Figure 3F-G**). The percentage of CD19^+^CD27^+^IgD^-^ class-switched B cells did not correlate with the age of donors of either sex, suggesting that the rise in hormone levels at puberty may be of greater significance than the age-related increase in the cumulative exposure to antigens (**Figure 3H-I**). Collectively, these results suggest that while sex chromosome-encoded genes play a potentially dominant role in the regulation of the B cell transcriptome, the presence of female sex hormones is needed to increase CSR in B cells.

**Figure 3:**
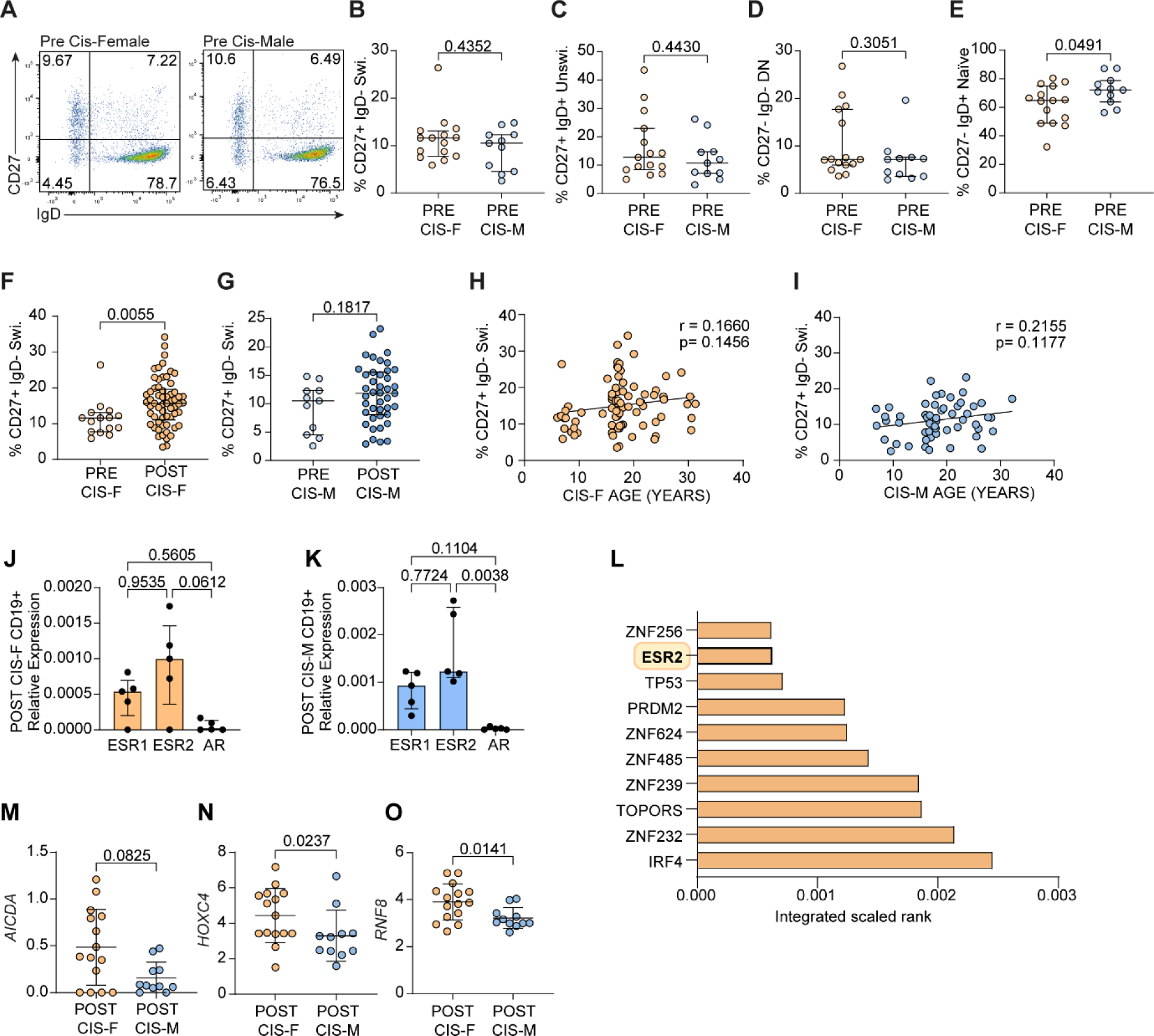
Oestrogen is associated with increased B cell class-switching in healthy cisgender females compared to age-matched males. (**A**) Representative flow cytometry plots and (**B-E**) summary dot plots comparing healthy, pre-pubertal cisgender female (PRE CIS-F; n=15) and male (PRE CIS-M; n=11) relative percentages of (**B**) class-switched memory (Swi.; CD27+ IgD-) (**C**) Unswitched (Unswi.;CD27+ IgD+), (**D**) Double Negative (DN; CD27-IgD-) and (**E**) Naïve (CD27-IgD+) B cells. (**F-G**) Comparison of class-switched B cell percentages in pre-pubertal versus post-pubertal (**F**) cisgender females (PRE CIS-F n=15; POST CIS-F n=61) and (**G**) cisgender males (PRE CIS-M n=11; POST CIS-M n=42). (**H-I**) Pearson test for correlation between age (years) and percentage of IgD-CD27+ switched B cells in cisgender females (**H**; n=78) and cisgender males (**I**; n=54). **(J-K)** Expression of hormone receptors in isolated CD19+ cells from post-pubertal cisgender female (POST CIS-F; n=5) and cisgender male (POST CIS-M; n=5) healthy donors, relative to expression of housekeeping gene RPLP0, measured by qPCR (Oestrogen Receptor Alpha -ESR1; Oestrogen Receptor Beta -ESR2 and Androgen Receptor -AR). (**L**) Top ten transcription factors whose putative transcriptional targets are most similar to identified class-switching gene set. Integrated score takes into account results from all ChEA3 transcription factor target libraries, with lower scores indicating higher relevance to the transcription factor. (**M-O**) Comparison of gene counts for AICDA, HOXC4 and RNF8 (key class-switching genes) from healthy POST CIS-F versus POST CIS-M B cell RNA sequencing. Mann-Whitney U-test (B-G & M-O) or Kruskal Wallis with Dunn’s test (J-K) with Median +IQR shown.

### Oestrogen has the capacity to directly modulate the expression of key transcriptional regulators of class-switch recombination

To understand the potential direct effect of oestrogen in regulating B cell CSR, we assessed the expression of sex hormone receptors (*ESR1*; Oestrogen Receptor alpha, *ESR2*; Oestrogen Receptor beta and *AR*; Androgen Receptor) in total B cells isolated from post-pubertal cisgender healthy females and males (**Figure 3J-K**). In both sexes, both *ESR1* and *ESR2* were expressed in B cells, with minimal expression of *AR*. To further this analysis, a set of genes known to regulate B cell class-switching (**Supplementary Table 2**) were assessed for putative transcriptional factor binding sites by comparing this class-switching gene set to ChEA3 libraries of transcription factor (TF) target gene sets. Of the 1,632 TFs included, *ESR2* was the 2^nd^ highest ranked TF (**Figure 3L**), demonstrating a direct ability for *ESR2*, but not *ESR1*, to regulate the expression of these genes. RNAseq gene counts of three class-switching genes (*AICDA, HOXC4, RNF8)* also show a strong trend toward an increase in cis-female compared to cis-male B cells (**Figure 3M-O**. **Supplementary Table 2** lists p-values for count comparisons of the other genes in this set, as well as their main function/s). Interestingly, studies from murine B cells have also demonstrated that B cells express oestrogen receptors, and that oestrogen can regulate expression of *AICDA* via *HOXC4* (Mai *et al*., 2010a).

### The impact of endogenous oestrogens on B cell class-switching is dependent on sex chromosomal complement

Together, our data suggest that both sex hormones and sex chromosomes play a role in controlling the heightened levels of class-switched memory B cells observed in cis-females, but did not directly address the relative contributions of both. To investigate this, we utilised a unique, gender-inclusive cohort which, due to modulation of sex hormone availability on both XX and XY chromosomal backgrounds, provides a novel opportunity to separate these contributing elements *in vivo*. Full explanations of the gender-affirming hormone therapy (GAHT) pathways are detailed in the Methods section and **Supplementary Figure 2**. Briefly, both transgender males (trans-males, XX karyotype, registered female at birth) and transgender females (trans-females, XY karyotype, registered male at birth) were commenced on ‘puberty blocker’ treatment, suppressing oestrogen/progesterone and testosterone production. Some trans-males then went on to take gender-affirming testosterone treatment, while some trans-females went on to take gender-affirming oestradiol (no additional progesterone or anti-androgen medication with progestogenic characteristics was administered).

We first compared the frequency of CD19^+^CD27^+^IgD^-^ B cells between post-pubertal cis-females, and their karyotype-counterparts, transgender males (trans-males) who had received either puberty blockers alone (which blocks the production of natal oestrogen/progesterone; referred to as *TRANS M+BLK* in figures) or puberty blockers with additional gender-affirming testosterone (referred to as *TRANS-M+T* in figures; **Figure 4A-E**). This analysis demonstrated that transgender males have a reduced frequency of CD19^+^CD27^+^IgD^-^ class-switched memory B cells and IgG^+^ B cells when compared to cisgender females *ex vivo*. Importantly, this decrease was observed following the blockade of oestrogen/progesterone using puberty blocker treatment, with gender-affirming testosterone treatment having no further effect on class-switched B cell frequencies (**Figure 4A-E** and **Supplementary Figure 1**).

**Figure 4:**
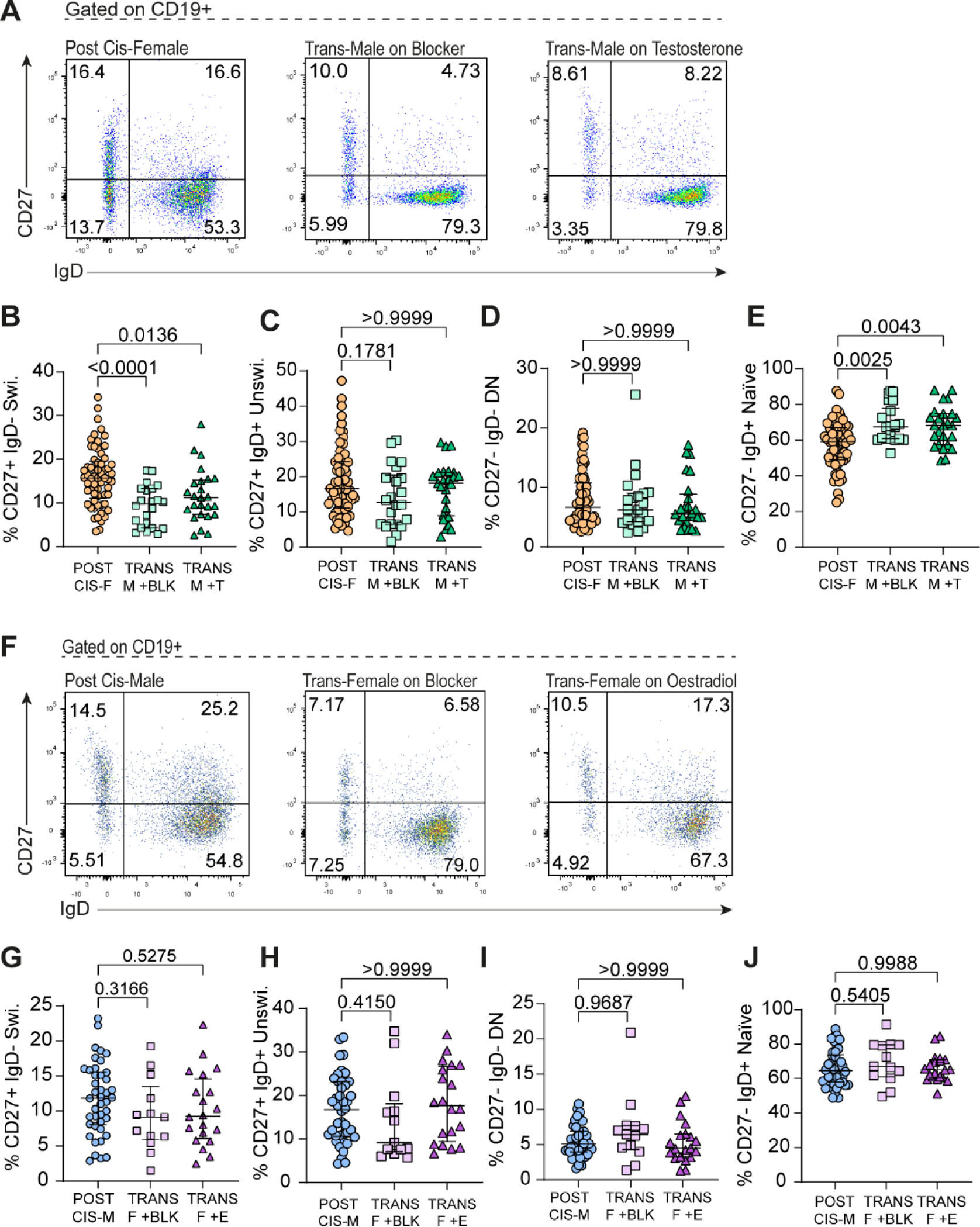
Differential impact of hormones on B cell class-switching dependent upon sex chromosomal complement. (**A**) Representative flow cytometry plots and (**B-E**) scatter plots showing relative percentages of (**B**) Class-switched memory (CD27+ IgD-), (**C**) Unswitched (CD27+ IgD+), (**D**) Double Negative (CD27-IgD-) and (**E**) Naïve (CD27-IgD+) B cells in post-pubertal cisgender females (POST CIS-F; n=61) compared to trans-males on GnRHa puberty hormone blockers (TRANS-M BLK; n=23) +/- gender-affirming testosterone treatment (TRANS-M + T; n=24). (**F**) Representative flow cytometry plots and (**G-J**) scatter plots showing relative percentages of (**G**) Class-switched memory (CD27+ IgD-), (**H**) Unswitched (CD27+ IgD+), (**I**) Double Negative (CD27-IgD-) and (**J**) Naïve (CD27-IgD+) B cells in post-pubertal cisgender males (POST CIS-M; n=42) compared to transgender females on puberty hormone blockers (TRANS-F BLK; n=15) +/- gender-affirming oestradiol treatment (TRANS-F + E; n=21). Ordinary one-way Analysis of Variance ANOVA with Tukey’s test and Mean +SD shown (B, G, J), or Kruskal Wallis test with Dunn’s post-hoc test and Median +IQR shown (C-E & H-I), as appropriate to distribution. Following correction for multiple testing, adjusted significance determined as p<0.01.

Conversely, when assessing the levels of class-switched B cells in transgender females in receipt of gender-affirming blockers (which block production of natal testosterone; *TRANS-F +BLK)* or additional oestradiol treatment (referred to as *TRANS-F +E* in figures) compared to post-pubertal cis-males, we found that there was no difference in both CD19^+^CD27^+^IgD^-^ or IgG^+^ class-switched B cell populations when compared to their chromosomal-counterparts, cisgender males (**Figure 4F-J**). Collectively, these results demonstrate that oestrogen is associated with increased levels of class-switched B cells *ex vivo* exclusively in individuals with an XX chromosomal background. This finding is of pertinence as there is a paucity of good-quality literature on transgender health and long-term outcomes for transgender populations, and demonstrates that hormonal transition may lead to distinct immunobiological gender phenotypes. This observation may be of growing importance as we approach the era of precision medicine.

Our study represents the first in-depth study of B cell immunobiology in cisgender and transgender individuals, providing a unique human-only insight into the mechanisms regulating B cell class-switching. Due to the clinical pathways by which these individuals are recruited there are some limitations to our study. As post-pubertal cisgender people are not recruited through a clinical pathway, we are unable to obtain total lymphocyte counts from these donors, meaning our data is based on cellular frequency rather than absolute numbers. In addition, trans-females do not receive progesterone or additional specific anti-androgen treatment as part of gender-affirming care (Milionis, Ilias and Koukkou, 2022), and our study does not rule out a possible additional role for physiological progesterone in the regulation of B cell class-switching, which has been suggested by limited murine literature (Pauklin and Petersen-Mahrt, 2009; Hughes and Choubey, 2014; Wong, Agrawal and Hughes, 2015). However, the progesterone receptor (PGR) was ranked only 321^st^ out of the 1632 transcription factors assessed for CSR gene binding sites, suggesting that oestrogens may be the dominant hormones influencing this process specifically. Future human studies are needed to address this and additional mechanistic exploration of how each hormone interacts with the class-switching process is warranted.

Despite these limitations, in this study we demonstrate an important complementary role for sex hormones and sex chromosomes in regulating the frequency of class-switched B cells. Our key observations include that only post-pubertal, but not pre-pubertal, cis-females have more class-switched B cells compared to cis-males. This shows that while sex-chromosomes drive most of the sex differences in B cell transcriptomes, this is insufficient to alter the class-switching element of B cell immunobiology. Further analysis demonstrates that oestrogen has a potentially direct role in regulating B cell class-switching as B cells express high levels of *ESR2* and genes regulating B cell class-switched are enriched for *ESR2* binding sites. Using a unique cohort of transgender individuals, we finally demonstrate that oestrogen works in tandem with an XX chromosomal background to increase the frequency of class-switched B cells. Blockade of oestrogen on an XX chromosomal background reduces the frequency of class-switched B cells, whilst addition of oestrogen on an XY chromosomal background has no impact on the levels of class-switched B cells. Thus, sex hormones differentially regulate B cell class switching in the presence of different sex chromosomes, suggesting that cisgender and transgender individuals have distinct immune phenotypes. This study adds to the growing evidence that sex and gender are critical factors to be considered in immunological studies. It also demonstrates that improvements in diversity and inclusion practises within medical research will not only advance our scientific understanding of sex-biases in disease outcomes, but potentially shed light on novel strategies for personally tailored healthcare.

## Acknowledgements

We would like to thank all the individuals who generously donated their blood and data to this study. Many thanks to Mr Jamie Evans of the UCL flow cytometry core facility in the Division of Medicine, and to Dr Diana Matei and Ms Persephone Jenkins (UCL Division of Medicine) for proofreading. We thank all members of the *Rheum Shared Seq* (RSS) study group who contributed to sample processing, metadata, data generation and curation, as well as analysis pipelines and data normalisation. We would also like to thank Professor Chris Wallace and Dr Wei-Yu Lin for supporting pipeline development for RNAseq analysis.

This work was funded by a Versus Arthritis PhD studentship to CC (22203) for HP. It was further supported by and hosted by the Centre for Adolescent Rheumatology Versus Arthritis at UCL, UCLH, and GOSH which is supported by Centre of Excellence grants from Versus Arthritis (21593 and 20164) and Great Ormond Street Hospital Children’s Charity awarded to LW. CC is supported by a grant from UCLH Biomedical Research Centre (BRC4/III/CC). ECR and NMDG were supported by a Medical Research Foundation ‘Lupus’ Fellowship awarded to ECR (MRF-057-0001-RG-ROSS-C0797). ECR is also supported by a Senior Fellowship from the Kennedy Trust for Rheumatology Research (KENN 21 22 09) and a FOREUM research career grant (094). LW and MK are supported by grants to the CLUSTER consortium, from the Medical Research Council (MRC) [MR/R013926/1], Versus Arthritis [Grant: 22084], Great Ormond Street Hospital Children’s Charity [VS0518], and Olivia’s Vision. LW and CD are also supported by the National Institute for Health and care Research (NIHR) Great Ormond Street Hospital Biomedical Research Centre. The views expressed are those of the author(s) and not necessarily those of the NHS, the NIHR or the Department of Health.

## Author contributions

HP, ECR, ECJ and CC conceived the project. HP, ECR, and CC designed experiments. HP, AR, NMDG, RR, MK, GAR and LMG performed experiments. HP, AR, CTD and RR analysed data. GB provided essential patients samples, clinical analysis and endocrinology expertise. HP and ECR co-wrote the manuscript. ECR, ECJ, LW, CD and CC obtained funding. CC critically reviewed the manuscript. All authors read and approved the final version of the manuscript.

## Methods

### Human participants

Full demographic information for participants is shown in **Table 1**. Post-pubertal cisgender healthy controls were recruited from the community, while pre-pubertal cisgender controls were recruited from otherwise healthy children undergoing elective corrective dental or urological surgeries at University College London Hospital (UCLH). ‘Post-puberty’ was self-reported by participants using a visual questionnaire as ‘Tanner stage 4-5’, and ‘pre-puberty’ was parentally reported as ‘Tanner stage 1’. Participants were screened for any history of autoimmune disease, relevant endocrinological conditions, serious or current infections and vaccination within the 3 weeks prior to blood donation.

Transgender healthy controls were recruited from the young persons’ Gender Identity Development Service (GIDS) at UCLH. Healthy, young transgender people were recruited to donate peripheral blood samples at determined intervals of gender-affirming hormone treatment (GAHT) regimens. Briefly, both transgender males (XX karyotype, registered female at birth) and transgender females (XY karyotype, registered male at birth) are commenced on Gonadotrophin Releasing Hormone analogues (GnRHa), commonly referred to as ‘puberty blockers’. Over-stimulation of the GnRH receptor leads to the eventual cessation of both oestrogen and subsequently progesterone, and testosterone production. After a minimum of 12+ months on GnRHa alone, those seeking further virilisation or feminisation and deemed Gillick competent (in the UK, persons under 16 years can consent to their own medical treatment if they’re felt to have suitable intelligence, competence and understanding to fully appreciate what’s involved in the treatment (Griffith, 2016)) may receive additional testosterone (trans-males) or oestradiol (trans-females), respectively, using gradually increasing dosages designed to mimic natal puberty. Samples are obtained when the young people have been on blockers (alone) for a minimum of four months, and then again when they have been on additional GAHT for a further minimum four months. Blocker treatment is maintained until natal oestrogen/testosterone levels are sufficiently suppressed by GAHT (and/or later gonadectomy in adulthood). This treatment pathway is also outlined in **Supplementary Figure 2**.

Of the trans-female participants, all were in receipt of GnRHa “puberty blocker” treatment for at least six months, and 20 had been additionally taking gender-affirming oestradiol (oral tablet or transdermal patch) for a further 4.5 months or more. Of the trans-male participants, all were also in receipt of at least six months of GnRHa blocker treatment, with 25 on additional gender-affirming testosterone (intramuscular injection or transdermal gel) for a further four months or more. 7 trans-males had acquired sufficient testosterone trough levels to dispense with GnRHa (required only until testosterone levels alone will sufficiently suppress natal oestrogen). All individuals in receipt of GnRHa blocker +/-gender-affirming hormones are encouraged to take calcium and vitamin D supplementation. 78/80 participants were deemed to have completed pubertal development as their birth-registered sex (Tanner Stage 4-5; based on both self-reported and clinician-confirmed tanner stage questionnaire) prior to commencement of any treatment, and 2/80 (both trans-females) were assessed as late Tanner stage 3.

Informed consent was taken from all volunteers aged 16 years or over. For those aged 6-15, or any person deemed not to have capacity; parental/guardian consent was taken (alongside assent from the volunteer themselves, where possible). All participants were recruited under North Harrow ethics committee approval REC11/0101. Basic health, demographic and puberty Tanner staging questionnaires were completed by/for all participants, and anonymised data was stored in compliance with data protection laws.

Venous whole blood was collected in heparin-coated vacutainers. In those under 18 years this was 2mL of blood per kg of weight, not exceeding a total of 20mL per sample. In those aged 18+, 20-60mL was taken.

### Peripheral blood mononuclear cell isolation and serum processing

Within two hours of sample collection, blood was diluted 1:1 with RPMI-1640 (Roswell Park Memorial Institute) medium containing L-glutamine and sodium bicarbonate (NaHCO3; Sigma-Aldrich) supplemented with 100 IU/μg/ml penicillin/streptomycin (Sigma-Aldrich) and 10% heat-inactivated foetal bovine serum (FBS; Gibco). SepMate™ tubes (Stemcell) were layered with Ficoll-Paque Plus (GE Healthcare), followed by the diluted sample, and then centrifuged at 1200g for 10 minutes. After further centrifugation of the top layer, pelleted isolated peripheral blood mononuclear cells (PBMC) were resuspended and counted using a haemocytometer, before being frozen in freezing media consisting of FBS with 10% dimethyl sulfoxide (DMSO; Sigma). Nalgene ‘Mr Frosty’ containers were used for controlled freezing at −80 °C, and samples were then transferred to liquid nitrogen storage.

### Flow cytometry

PBMC samples were thawed and washed once in warm RPMI-1640 media with 10% FBS (“cRPMI”; complete RPMI). Cells were counted with a haemocytometer. Thawed cells were plated at either 250k, 500k or 1 million cells per well (depending on the panel) in a 96 well plate and centrifuged at 500g for 5 minutes to pellet. Following resuspension in Phosphate Buffered Saline (PBS), they were stained with fixable blue dead cell stain (1:500) (ThermoFisher, L23105) for 15 minutes at room temperature in foil. After washing, cells receiving the immunoglobulin panel stain *(CD19-BV510 [HIB19], CD27-BV711 [O323], IgD-BV421 [IA6-2], IgM-FITC [MHM-88], IgA-APC [IS11 8E10], IgE-PE [IGE21], IgG-BUV737 [G18-145])* were first resuspended in FACS buffer (1x Dulbecco’s Phosphate Buffered Saline (DPBS) without calcium or magnesium (GE Healthcare Hyclone SH30028.02) + 5 ml FBS + 2mM Ethylenediaminetetraacetic acid [EDTA]) with Human TruStain FcX™ Fc Receptor Blocking Solution (1:20; BioLegend) for 15 minutes at room temperature in foil. Without washing, cells were spun down to pellet. For the surface staining, cells were incubated in foil for 30 minutes at room temperature with the relevant antibody panel *(Panel 1-B cells and monocytes: BADCA2-PE [201A], CD56-FITC [5.1H11], CD16-APC [3G8], CD123-PE/Cy7 [6H6], HLA-DR-PERCPCy5.5 [L243], CD24-APC/Cy7 [ML5], CD14-AF700 [63D3], IgD-BV421 [IA6-2], CD19-BV510 [HIB19], CD38-BV605 [HIT2], CD11c-BV785 [3.9], CD27-BV711 [0323], CD3-FITC [HIT3a], CD66b-FITC [G10F5]; Panel 2-T cells & Natural Killer cells: CCR7-PE [G043H7], TCRγδ-FITC [11F2], CD25-APC [BC96], CD56-PE/Cy7 [MEM-188], CD45RO-PERCPCy5.5 [UCHL1], CD4-APC/Cy7 [OKT4], CD8-AF700 [SK1], CXCR3-BV421 [G025H7], CCR6-BV510 [G034E3], CD3-BV605 [OKT3], CXCR5-BV785 [J252D4], CD127-BV711 [A019D5], CD45RA-PEDAZZLE [HI100]),* diluted in FACS buffer.

After washing, surface stain-only panels were fixed with 2% Paraformaldehyde Fixative (PFA) for 15 minutes in foil at room temperature, before washing and resuspension in FACS buffer and storage at 4°C. Panels requiring intracellular staining were fixed using the eBioscience Foxp3 staining kit (00-5523-00) Fix/Perm, for 15 minutes in the dark, before washing and staining with the relevant intracellular antibody mix/es for 40 minutes in the dark, followed by washing with perm buffer and resuspension in FACS buffer for 4°C storage.

All samples were run the following day on a BD LSRII flow cytometer, with as many cellular events as possible recorded per sample. Data was analysed using Flowjo v10, with a minimum of 100 events in the parent gate of any population used in analysis. Gating strategies were determined using Fluorescence Minus Ones (FMOs).

### Fluorescence-activated cell sorting (FACS)

Samples were thawed and counted as above, before staining with Zombie NIR™ Fixable dye (BioLegend) in cold DPBS on ice for 20 minutes. After washing, samples were stained with the following antibody panel: *CD4-BUV395 [SK3], CD8-BV785 [RPA-T8], CD19-AF488 [HIB19], CD14-PE/Cy7 [M5E2]*, in cold FACS buffer, on ice for 20 minutes. They were then washed and filtered into filter-capped FACS tubes in cold FACS buffer and sorted into cold DPBS using a BD FACSAria flow cytometry cell sorter.

### RNA Extraction

RNA was extracted and isolated from sorted CD4/CD8/CD14/CD19+ cells, and from MCF7 cells as a control, known to be positive for hormone receptors (Horwitz et al., 1975). The PicoPure™ RNA Isolation Kit (ThermoFisher) was used, following the manufacturer’s guidelines. The RNA columns were additionally incubated with RNase-free DNase (Quiagen) at room temperature for 15 minutes during this process. Eluted RNA was quantified using a NanoDrop spectrophotometer. *Reverse transcription-polymerase chain reaction (RT-PCR) for cDNA Synthesis* cDNA was synthesised from RNA following the manufacturer’s instructions of the qScript cDNA SuperMix (Quantabio). The GeneAmp PCR System 9700 Thermocycler was used with the following settings: ***5 minutes at 25°C, 30 minutes at 42°C, 5 minutes at 85°C, Hold at 4°C*.**

### qPCR

Applied Biosystems™ TaqMan™ Fast Advanced Master Mix (ThermoFisher) was added to cDNA for fluorescence quantification on an *AriaMx Real-time qPCR System* (Agilent Technologies) using the following thermal profile: **UDG (DNA)** 2min 50°C, **Polymerase Activation Hold** 95°C 2min, **Denature** 95°C 1 second, **Anneal/Extend** 60°c 20 seconds.

TaqMan™ probes for Androgen Receptor (Hs00171172_m1, 4331182), Oestrogen Receptor alpha (“*ESR1*”- Hs00174860_m1, 4331182) Oestrogen Receptor beta (“*ESR2*”- Hs00230957_m1, 4331182), Activation-Induced Cytidine Deaminase (“*AICDA”-* Hs00757808_m1, 4331182) and housekeeping gene RPLP0 (60S acidic ribosomal protein P0) (4333761T) were purchased from ThermoFisher and used at the manufacturer’s recommended concentrations for each reaction. *Agilent Aria 1.7* software was used to prepare the data, and Microsoft Excel 2008 was used for analysis. Data was normalised to the reference gene RPLP0. The Delta-Ct (Livak & Schmittgen, 2001) method of quantification was used to ascertain relative gene expression of the target genes.

### RNA sequencing

RNA sequencing was performed by UCL Genomics (UCLG). Briefly, libraries were prepped with either the *KAPA mRNA HyperPrep Kit* or the *TruSeq Stranded mRNA Library prep.* Paired-reads mapping was performed using *STAR aligner* (Dobin *et al*., 2013) (Ref genome: GRCh38) and aligned reads were summarised with *featureCounts.* Quality control was conducted on the bulk RNA-seq reads count-table obtained from *featureCounts* to ensure data quality. Samples with fewer than 5 million reads were excluded, as these were deemed insufficient for reliable analysis.

Principal component analysis (PCA) was subsequently employed to identify outliers within each cell population. It was assumed that samples from the same cell population would cluster together. PCA was performed on the filtered count-table, and the first two principal components (PC1 and PC2) were plotted to visualize sample clustering. For each cell population, the centre and standard deviations of the clusters were calculated. Samples deviating more than 4 standard deviations from the cluster centre were identified as outliers and removed. This process ensured that only high-quality, reliable samples were retained for downstream analysis. Detailed RNA sequencing data analysis workflow describing quality control, alignment, and generation of raw transcript counts can be found in the following GitHub repository: https://github.com/WedderburnLab/RNAseq-Pipeline. Read counts were kit-corrected with *CombatSeq* (Zhang, Parmigiani and Johnson, 2020) prior to differential gene expression analysis.

Statistical analysis and visualisation of transcriptional data was performed using R software and Bioconductor packages (Gentleman *et al*., 2004) including DESeq2 (Anders and Huber, 2010; Love, Huber and Anders, 2014) and EnhancedVolcano (Blighe K, Rana S and Lewis M, 2022).

Genes for which 50%+ of the control group had counts of 10 or less were excluded from the analysis. DESeq2 in R was used to carry out differential gene expression (DEG) analysis, between two selected groups to calculate the fold changes and the adjusted p-value (padj) for significance of differences (adjusted for multiple testing using Benjamini-Hochberg’s false discovery rate (FDR) method). Both age of donor and batch that the sample was sent in were controlled for within the analysis model. A padj cut-off of <0.05 was applied using Microsoft Excel, to generate DEG lists for each comparison. Those with a positive fold change were considered to be upregulated, and those with a negative fold change to be downregulated. Detailed RNA sequencing data analysis workflow describing quality control, alignment, and generation of raw transcript counts can be found in the following GitHub repository: https://github.com/WedderburnLab/RNAseq-Pipeline

Pathway enrichment analysis (PEA) was conducted with separate lists of up- and downregulated genes for each comparison, using Metascape (Zhou *et al*., 2019) to reveal significantly up/downregulated ‘GO: Biological Processes’ pathways (Ashburner *et al*., 2000; Aleksander *et al*., 2023) within each comparison. Pathways were ranked by p-value into bar charts, shown alongside the ‘enrichment score’ (the ratio of the proportion of genes in the list that were associated with the enriched term to the proportion of genes in the genome that are associated with the enriched term). For reduction of redundancy, the program clusters similar enrichment terms, with the most significant term of each cluster represented on the bar chart. DEG lists were further summarised in heatmaps (created with *Clustvis* (Metsalu and Vilo, 2015), using Euclidian distance and the average method for clustering of rows) for each comparison, demonstrating the clustering of each group by expression of the listed genes. Targeted gene analysis using the gene set detailed in **Supplementary Table 2** was carried out using count data (transcripts per million, TPM) for each sample, and analysed using GraphPad Prism. Briefly, the lists of genes from the ‘GO Biological Pathways’ ontology pathway ‘GO:0045190-Isotype Switching’, alongside the lists from ‘GO:0045191-regulation of isotype switching’ and ‘GO:0048291-regulation of isotype switching to IgG’ pathways were cross-examined, and duplicates removed. Also removed were any genes deemed not to be expressed on human B cells, according to The Human Protein Atlas (*The human protein atlas*, 2000), and the accompanying supporting literature from the GO website (Ashburner *et al*., 2000; Aleksander *et al*., 2023). Genes *HOXC4* and *BACH2* were included, given their prominence in the literature on sex differences in CSR to date. To assess enrichment of transcription factor binding sites in these genes, genes were inputted into the ChEA3 online tool (Keenan et al., 2019).

### Statistical Analysis

Statistical analysis was carried out using *GraphPad Prism 10*. Unpaired t-tests, Mann-Whitney U-tests, one-way ANOVA with Tukey’s test, or Kruskal-Wallis test with Dunn’s test (dependent upon normality of each dataset and number of groups being compared) were used to determine differences between groups. P-values adjusted for multiple testing were calculated by dividing 0.05 by the number of tests performed.

