## Supplementary Materials for "Oestrogen influences B cell class-switching in individuals with an XX sex chromosome complement"

### Supplementary Tables and Figures

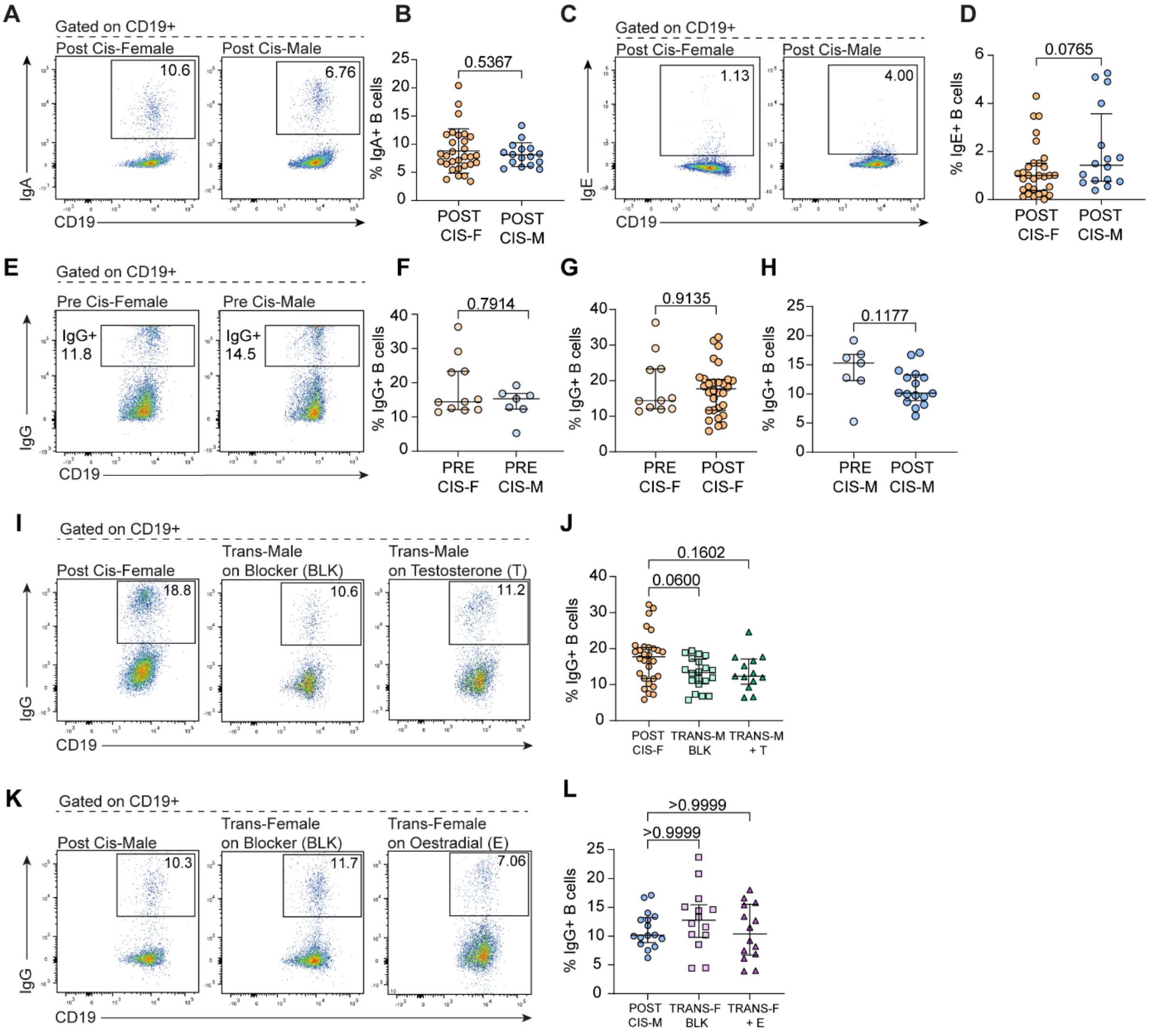


***Supplementary Figure 1: Class-Switched B cells in natal and hormonally-induced puberty** (**A & C**) Representative flow cytometry plots and (**B & D**) scatter plots showing relative percentages of (**A-B**) IgA+ B cells and (**C-D**) IgE+ B cells in post-pubertal cisgender females (POST CIS-F, n=) versus post-pubertal cisgender males (POST CIS-M, n=). (**E**) Representative flow cytometry plots and (**F-H**) scatter plots comparing relative percentages of IgG+ B cells between (**F**) pre-pubertal cisgender females (PRE CIS-F, n=11) and pre-pubertal cisgender males (PRE CIS-M, n=7), (**G**) pre cis-F and post-pubertal cisgender females (POST CIS-F, n=30) and (**H**) pre cis-m and post-pubertal cisgender males (POST CIS-M, n=17). (**I**) Representative flow cytometry plots and (**J**) scatter plot comparing relative percentages of IgG+ B cells between post cis-F and transgender males on puberty blocker (TRANS-M BLK, n=20) and transgender males on gender-affirming testosterone (TRANS-M +T, n=13). (**K**) Representative flow cytometry plots and (**L**) scatter plot comparing relative percentages of IgG+ B cells between post cis-M and transgender females on puberty blockers (TRANS-F BLK, n=14) and transgender females on gender-affirming oestradiol (TRANS-F +E, n=14). Unpaired t-test with Mean +SD (B), Mann-Whitney U-test with Median +IQR (D, F-H) or Kruskal Wallis test with Dunn's post-hoc test and Median +IQR (J & L) as appropriate to distribution of data.*

| Down in POST CF vs. POST CM | Chromosome | Up in POST CF vs. POST CM | Chromosome |
|---|---|---|---|
| CHIT1 | 1 | EIF1AXP1 | 1 |
| ZAP70 | 2 | MARCHF7 | 2 |
| PRF1 | 10 | RGPD2 | 2 |
| AC136475.3 | 11 | RGPD1 | 2 |
| PXN | 12 | ZNF141 | 4 |
| HRK | 12 | BNIP3P41 | 4 |
| GZMH | 14 | TMEM106B | 7 |
| GPR68 | 14 | MS4A7 | 11 |
| CCL5 | 17 | IRAK3 | 12 |
| AC254562.3 | 22 | EIF2S3B | 12 |
| KDM5D | Y | WDR89 | 14 |
| DDX3Y | Y | ZNF681 | 19 |
| ZFY | Y | ZNF181 | 19 |
| PRKY | Y | ZNF480 | 19 |
| USP9Y | Y | ZFX | X |
| RPS4Y1 | Y | PNPLA4 | X |
| TXLNGY | Y | SMC1A | X |
| TTTY14 | Y | TXLNG | X |
| UTY | Y | USP9X | X |
| EIF1AY | Y | PUDP | X |
| BCORP1 | Y | EIF2S3 | X |
| AC010086.1 | Y | KDM6A | X |
| TTTY10 | Y | ZRSR2 | X |
| LINC00278 | Y | EIF1AX | X |
| ANOS2P | Y | PRKX | X |
| AC010889.1 | Y | MAP7D2 | X |
| RPS4Y2 | Y | CXorf38 | X |
| AC244213.1 | Y | ERCC6L | X |
| | | TRAPPC2 | X |
| | | RPS4X | X |
| | | DDX3X | X |
| | | INE1 | X |
| | | JPX | X |
| | | XIST | X |
| | | TSIX | X |
| | | BX890604.1 | X |

***Supplementary Table 1: Significantly up- or downregulated genes in B cells from post-pubertal cisgender females (POST CF) vs. post-pubertal cisgender males (POST CM)***

| Gene | Significance to Class-Switch Recombination | Cis-F vs. Cis-M gene count p-value |
|---|---|---|
| *AICDA* | Activation-induced cytidine deaminase. Critical enzyme for the DNA mutations necessary for CSR to occur (Muramatsu *et al.*, 2000). | 0.0825 |
| *ATAD5* | ATPase Family AAA Domain Containing 5. Responsible for unloading proliferating cell nuclear antigen from newly synthesised DNA. In knock-out mice, AID expression, CSR and B cell division are all decreased (Zanotti *et al.*, 2015). | 0.6098 |
| *BATF* | Basic Leucine Zipper ATF-Like Transcription Factor. Controls AID expression (Ise *et al.*, 2011). | 0.2171 |
| *BCL6* | B-cell lymphoma 6. Transcriptional repressor necessary for formation of germinal centres, preventing B cells from becoming short-lived plasma cells (Alinikula *et al.*, 2011). BCL-/- mice exhibit increased CSR to IgE and inflammatory symptoms (Harris *et al.*, 1999) as it binds to the E-germline transcription promoter (Audzevich *et al.*, 2013). Depends on **STAT6** (Haase *et al.*, 2020)**.** | 0.0705 |
| *BACH2* | A transcriptional regulator which limits B cell differentiation into plasma cells until after AID expression, when CSR and SHM have consequently taken place. Compromised expression in SLE. Bach2-/- mice have enhanced extra-follicular CSR & IgG+ autoantibodies (Jang *et al.*, 2019). | 0.9381 |
| *CCR6* | Significant role in switch to IgA and gut immunity (Lin, Ip and Liao, 2017). | 0.1592 |
| *ERCC1* | Excision Repair 1. ERCC1-XPF shown to be essential component of DNA repair pathway, interacts with **MSH2** in processing or repairing DNA lesions in S regions in CSR (Schrader *et al.*, 2004). | 0.9787 |
| *EXO1* | Exonuclease 1. Exo-/- mice had decreased CSR (Bardwell *et al.*, 2004; Eccleston *et al.*, 2011). Acts with **MSH2** and **MLH1.** | 0.5065 |
| *EXOSC3* | Exosome Component 3. EXOSC3-deficient B cells cannot class switch (Pefanis *et al.*, 2014). | 0.1287 |
| *EXOSC6* | Exosome Component 6. Subunit of the exosome, which *may* target AICDA deaminase activity toward transcribed dsDNA substrates Basu et al., 2011). | 0.9777 |
| *HOXC4* | A transcription factor- binds directly to the *AICDA* gene promoter, thus potentiating CSR. Upregulated in SLE & lupus-prone mice. *Hoxc4* -/- mice showed a reduction in IgG2a autoantibodies and IgG kidney deposition (White *et al.*, 2011). Oestrogen has also been demonstrated to upregulate *Hoxc4*. In lupus-prone mice, elevated levels of class-switched autoantibodies and chromosomal translocations, induced in a Hoxc4-AID-dependent manner, suggest that oestrogen may play a part in AID dysregulation in SLE (Pauklin *et al.*, 2009; Mai *et al.*, 2010b). | 0.0237 |
| *LIG4* | DNA Ligase 4. NHEJ factor (Ghosh and Raghavan, 2021). Mutations in LIG4 lead to "*LIG4 syndrome*", where patients have less effective or defective V(D)J recombination. Some | 0.9957 |

| | | |
|---|---|---|
| | have a severe immunodeficiency (Altmann and Gennery, 2016). | |
| *MLH1* | MutL Homolog 1. Mismatch repair protein, thought to convert nicks to breaks, acts with **MSH2 & EXO1** (Eccleston *et al.*, 2011). | 0.1574 |
| *MSH2* | MutS Homolog 2. In mice, involved in DNA mismatch recognition. Msh2-/- show decreased IgG *in vitro* and *in vivo* (Ehrenstein, 1999). Interacts with **ERCC1.** | 0.2826 |
| *MSH6* | MutS Homolog 6. A mismatch repair protein (MMR), its phosphorylation regulates CSR via DYRK1A (Stoler-Barak *et al.*, 2023). | 0.1723 |
| *NBN* | Nubrin protein - role in double stranded break repair as part of Mre11/Rad50/Nibrin complex (Piątosa *et al.*, 2012) and recombination of Ig constant genes (Kracker et al., 2005)). | 0.0654 |
| *NFKBIZ* | IKBNS-/- B cells in mice have decreased proliferation and CSR to IgG3. Demonstrated role in TLR-medicated T-independent CSR (Touma *et al.*, 2011). | 0.0697 |
| *RNF168* | Ring Finger Protein 168. Double stranded breaks are repaired by 53BP-1-dependent process - **RNF8** & RNF186 recruit 53BP-1 to site of damage (Ramachandran *et al.*, 2010). | 0.7982 |
| *RNF8* | Ring Finger Protein 8. Key ubiquitination pathway mediator, integral to CSR (Ramachandran *et al.*, 2010). RNF8 deficient mice show impaired CSR (Li *et al.*, 2010) and RNF8 knock-out mice had defective CSR and impaired double stranded breaks (Santos et al., 2010) (Santos et al., 2010). | 0.0141 |
| *STAT6* | Signal Transducer And Activator Of Transcription 6. Crucial role in B cell development & CSR (Wang, Wang and Zha, 2021). In mice, pivotal role in IgG1 and IgE switch (Haase *et al.*, 2020). Linked pathway with **BCL6.** | 0.7325 |
| *SWAP70* | Switching B Cell Complex Subunit. SWAP70 knock-out mice exhibit decreased IgE & IgG1 production (Borggrefe *et al.*, 2001) with IgE production controlled via **STAT6/BCL6.** | 0.0862 |
| *UNG* | Controversial role in CSR (Yousif *et al.*, 2014). *UNG*-/- mice were deficient in CSR (Rada *et al.*, 2002), unclear mechanism. | 0.1878 |

***Supplementary Table 2: Class-switching gene set used in fig 3 L-O.*** *CSR- class-switch recombination; DNA - deoxyribonucleic acid; ATP- adenosine triphosphate; AID- activation-induced deaminase; SHM- somatic hypermutation; SLE- systemic lupus erythematosus; NHEJ- non-homologous end joining; Ig- immunoglobulin. Mann-Whitney U test used to obtain p-values.*

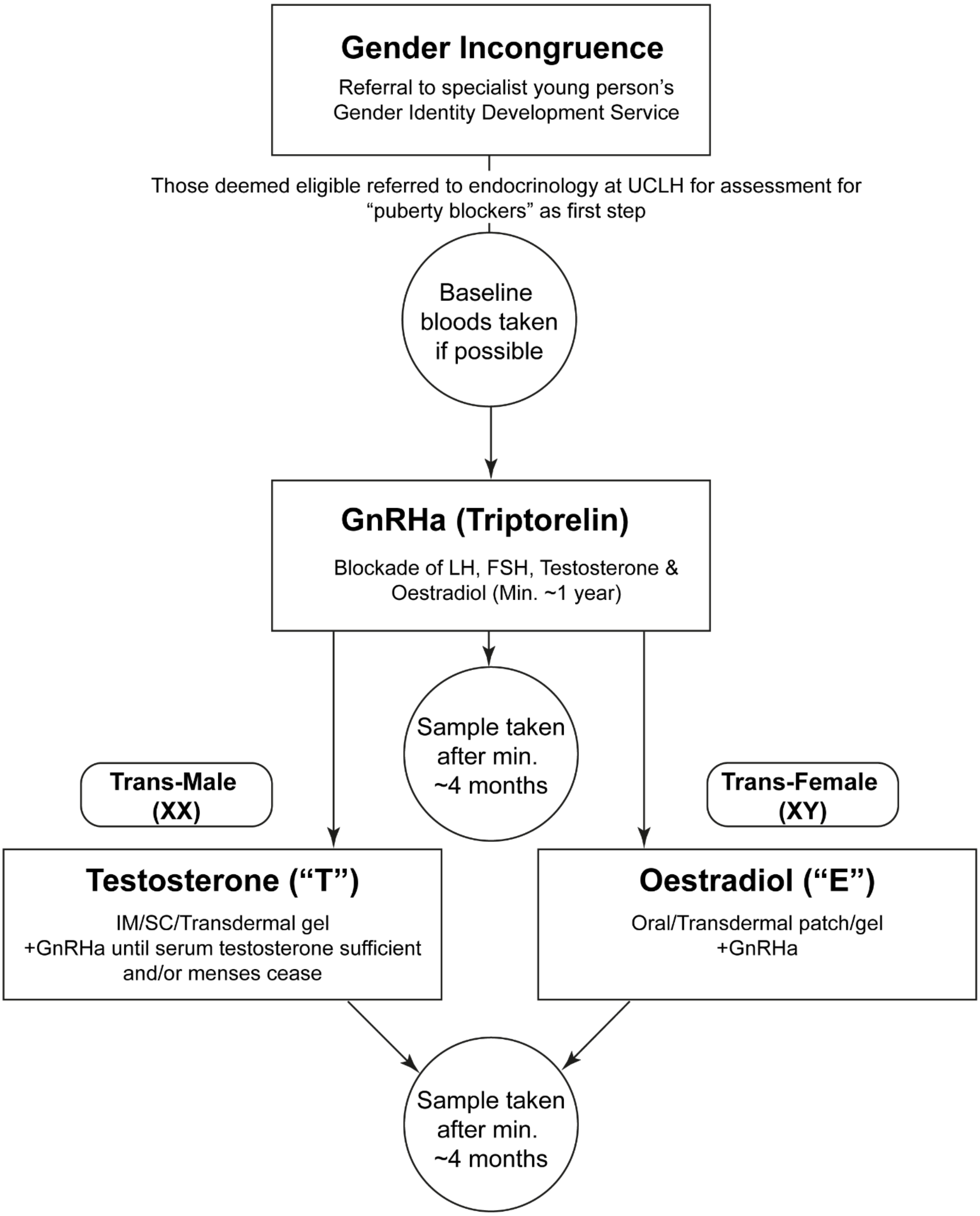


***Supplementary Figure 2: Gender Clinic Treatment Pathway and sample schedule*** *Treatment is prescribed on a case-by-case basis, based on individual country guidelines. This flowchart outlines the most commonly pursued routes by young persons referred to England's National Health Service (NHS) Gender Identity Development Service (GIDS).* GnRHa*: Gonadotropin releasing hormone analogues,* LH*: Luteinising hormone,* FSH*: Follicle stimulating hormone,* IM*: Intramuscular,* SC*: Subcutaneous. Summarised from guidelines outlined in Hembree et al, (2017).*
